# Polyploidization enhancing genetic recombination of the ancestral diploid genome in the evolution of hexaploid wheat

**DOI:** 10.1101/2020.02.21.958991

**Authors:** Hongshen Wan, Jun Li, Shengwei Ma, Qin Wang, Xinguo Zhu, Zehou Liu, Fan Yang, Manyu Yang, Jianmin Zheng, Shizhao Li, Jiangtao Luo, Wuyun Yang

**Affiliations:** Crop Research Institute, Sichuan Academy of Agricultural Sciences, Chengdu 610066, China; Key Laboratory of Wheat Biology and Genetic Improvement on Southwestern China (Ministry of Agriculture and Rural Areas), Chengdu 610066, China; Triticeae Research Institute, Sichuan Agricultural University, Chengdu 611130, China; The Applied Plant Genomics Lab, College of Agricultural Sciences, Nanjing Agricultural University, Nanjing 210095, China

**Author notes:** Correspondence: Crop Research Institute, Sichuan Academy of Agricultural Science Chengdu 610066, Sichuan, China. H. Wan and J. Li contributed equally to this article.

**Keywords:** Polyploidization, genetic recombination, diploid genome, enhancing evolution, synthetic hexaploid wheat

## Abstract

Allopolyploidy increases its evolutionary potential by fixing heterosis and the advantage of gene redundancy. Allelic combinations generated from genetic recombination potentially provide many variations to the selection pools for evolution. May there be any relationship between allopolyploidization and genetic recombination? To study the impact of polyploidy on genetic recombination, we selected wheat as a model and simulated its evolution pathway of allopolyploidy by developing synthetic hexaploid wheat. The change of homologous chromosome recombination were investigated on their diploid DD and tetraploid AABB genomes after their allohexaploidization, respectively. The genetic recombination of the ancestral diploid genome of *Aegilops tauschii* was enhanced significantly more than 2 folds after their hexaploidization. Hexaploidization enhancing genetic recombination of the ancestral diploid D genome was firstly reported to be a new way to increase evolutionary potential of wheat, which is beneficial for wheat to conquer their narrow origination of D genome, quickly spread and make it a major crop of the world. Finally, re-synthetizing hexaploid wheat using diverse *Ae. tauschii* species with tetraploid wheat can be considered as a pleiotropic strategy to speed adaptive evolution of bread wheat in breeding processes by increasing both gene allele types and genetic recombination variations.

## Introduction

Polyploidy occurs throughout the evolutionary history in many important agricultural crops (Leitch and Bennett, 1997; Udall and Wendel, 2006). It can be classified into autopolyploids such as potato (2n=4x=48) and allopolyploids such as oilseed rape (2n=4x=38, AACC), bread wheat (2n=6x=42, AABBDD), and synthetic hexaploid and octoploid *Triticale* (2n=6x=42, AABBRR and 2n=8x=56, AABBDDRR), based on the origins and levels of ploidy (Chen and Ni, 2006; Chen, 2007). Polyploidy takes its advantage on heterosis (in wheat: Li et al., 2014a; Li et al., 2014b) and gene redundancy (Comai, 2005), which might instead increase diversity, plasticity and adaptation of polyploids after overcoming the disadvantages of difficulties in the normal completion of meiosis (in wheat: Dubcovsky et al., 1995; Prieto et al., 2004) and genetic stability (Feldman et al., 1997; Shaked et al., 2001; Zhang et al., 2013). Heterosis causes polyploidy to be more vigorous than its diploid progenitors, whereas gene redundancy shields polyploidy from the deleterious effect of mutations, therefore enhancing its evolutionary potential (Comai, 2005; Paterson, 2005).

Wheat was domesticated about 10,000 years ago and has since spread worldwide to become one of the major crops. It originated by hybridization of tetraploid *Triticum turgidum* with diploid *Aegilops tauschii*. Among the tetraploid wheat, *T. turgidum* ssp. *dicoccum* was domesticated from the wild plant *T. turgidum* ssp. *dicoccoides* in the Fertile Crescent and the larger Near East about 10,000 years ago and spread over the next ∼2,000-3,000 years northwest across Anatolia and southeast to the southern Zagros mountains and to the northwest into Europe, southwest to the Nile Valley, southeast towards the Indus Valley, and northeast to Central Asia at later dates (Bar-Yosef and Meadow, 1995). Interestingly, the wild diploid species *Ae. tauschii* ssp. *strangulate* as the donor of the D genome (Nakai, 1979; Jaaska, 1980; Nishikawa et al., 1980; Lagudah et al., 1991; Lubbers et al., 1991; Dvorak et al., 1998, 2012; Wang et al., 2013) hybridizing with the free-threshing form of tetraploid wheat (Dvorak et al., 2012), which most probably happened in the south and west of the Caspian Sea about 9,000 years ago, made the hexaploid wheat being a major type of wheat in the next agricultural history, which accounting for about 95% of world wheat production whereas the tetraploid durum wheat (*T. durum*) only accounting for the other 5% (Dubcovsky and Dvorak, 2007).

*Ae. tauschii*, the D genome donor of the common wheat, is a widely distributed (van Slageren, 1994) and genetically diverse species (Lubbers et al., 1991; Dvorak et al., 1998; Schneider et al., 2008). A total of four morphological varieties have been found in nature, of which three are grouped into *Ae. tauschii* ssp. *tauschii* mostly called as ‘lineage 1’ (Mizuno et al.,2010; Wang et al., 2013), including var. *typica*, var. *anathera* and var. *meyeri* (Kihara and Tanaka, 1958), and the fourth is grouped into *Ae. tauschii* ssp. *strangulata* (Kihara and Tanaka, 1958), broadly related to ‘lineage 2’ (Mizuno et al.,2010; Wang et al., 2013)*. Ae. tauschii* worked as a gene repertoire for modern wheat adaptation (Jia et al., 2013; Li et al., 2014b), for their superiority in resistance to disease (Innes and Kerber, 1994; Cox et al., 1994; Yang et al., 2003; Miranda et al., 2006; Olson et al., 2013a, 2013b) and pest (Cox and Hatchett, 1994; Thompson and Haak, 1997; Zhu et al., 2005), tolerance to environmental stresses (Schachtman et al., 1991; Lan et al., 1997; Pritchard et al., 2002; Ryan et al., 2010; Sohail et al., 2011), and yield enhancement (Li et al., 2002; Huang et al., 2004; Kumar et al., 2007; Wan et al., 2015), all of which could be fixed as heterosis in hexaploid wheat theoretically.

However, the D genome of the first bread wheat is originated from only a small part of *Ae. tauschii* ssp. *strangulate* (Nakai, 1979; Jaaska, 1980; Nishikawa et al., 1980; Lagudah et al., 1991; Dvorak et al., 1998, 2012; Wang et al., 2013), and their genetic basis and diversity are rather narrow in modern common wheat. To broaden the D genome’s diversity for benefitting wheat breeding, scientists tried to transfer the interesting genes of *Ae. tauschii* to wheat by direct hybridization, and unfortunately got rare success as the cross between hexaploid wheat and *Ae. tauschii* were often sterile and only few genomic components of *Ae. tauschii* could be reserved even by the embryo rescue technique or backcrossing (Riley and Chapman, 1960; Raupp et al., 1983; Alonso and Kimber, 1984; Gill and Raupp, 1987). The creation of synthetic hexaploid wheat (SHW) by using tetraploid wheat crossing with *Ae. tauschii* with chromosome doubling makes the genetic transfers from *Ae. tauschii* to wheat more efficient and much easier (Mujeeb-Kazi et al., 1996), and lots of elite commercial wheat varieties derived from SHW have been released in the subsequent decades (Yang et al., 2009; Mujeeb-Kazi et al., 2013, Li et al., 2018). To some extent, the artificial polyploidization also enhanced the genetic variation and adaptive evolution of bread wheat in breeding processes (Li et al., 2014a).

Genetic recombination results in novel allele combinations and is the most important genetic phenomenon for generating variation in breeding populations. Genetic advance in a breeding program depends on selection of new recombinant individuals from inter-varietal or interspecific crosses. Genetic recombination can generate novel allele combinations for providing lots of genetic variation to the selection pools for evolution. And boosting genetic recombination would effectively speed up the combining of valuable traits from different parents in new elite varieties (Wijnker and de Jong, 2008). Allopolyploidization has been a driving force in plant evolution (Ozkan et al., 2001; Feldman and Levy, 2005), which has been focused primarily on by plant evolutionary genomicists in recent years. The commonly held view is that polyploidy takes its advantage on heterosis and gene redundancy, but the change of genetic recombination such as homologous recombination of polyploidy, which might be a a strong force driving some aspects of plant genome variability (Gaut et al., 2007), has not received adequate attention yet, especially in wheat evolution.

To study the impact of allopolyploidy on genetic recombination in wheat, we developed two sets of segregation populations to investigate the change of the genetic recombination frequency between homologous chromosomes of AB, and D genomes from tetraploid and diploid to hexaploid, respectively. With the help of molecular markers, the distribution of genome-wide recombination frequency and their changing pattern along entire chromosomes of A, B and D genomes were examined under diploid, tetraploid and hexaploid levels. The results explicitly declared that allohexaploidization enhanced the homologous recombination of the diploid donor genome of common wheat, suggesting that polyploidization could speed adaptive evolution of bread wheat by enhancing the genetic recombination of the ancestral diploid genome, and providing a theoretical support of re-synthetizing hexaploid wheat using different diploid donor genome and then making them crossing with each other to manipulate recombination in the creation of diversified genetic resources for wheat breeding.

## Materials and Methods

### Plant materials

Two *Ae. tauschii* accessions SQ665 and SQ783 (*Ae. tauschii* ssp. *tauschii* var. *typica*) and two *T. turgidum* cultivars Yuanwang (*T. turgidum* conv. *turgidum*) and Langdon (*T. turgidum* conv. *durum*) were used to generate three SHWs Langdon/SQ665 (LS665), Langdon/SQ783 (LS783) and Yuanwang/SQ783 (YS783). The two *Ae. tauschii* accessions were provided by Dr. A. Mujeeb-Kazi at the International Wheat and Maize Improvement Centre (CIMMYT), Mexico, in 1995. The teraploid wheat Yuanwang is a landrace of *T. turgidum* conv. *turgidum* collected in Sichuan, China. In this study, two tetraploid wheats were used as female parents, while two *A. tauschii* accessions were used as male parents. In their first selfed generation (S1) of LS665, LS783, and YS783, most of their offspring contained the euploid chromosome set (2n=42), which were karyotyped by Zhu (2017), using fluorescence in situ hybridization (FISH) with two repetitive DNA sequences *Oligo-pSc119.2* labeled with Alexa Fluor 488-5-dUTP (green coloration) and *Oligo-pTa535* with Texas Red-5-dCTP (red coloration) (synthesized by Invitrogen, Shanghai, China), which have successfully identified all 21 homologous chromosomes pairs of Chinese Spring (CS) (Tang et al. 2014). Among these offspring from S2, we selected individuals covering the whole A, B and D genome with 42 chromosomes and then created two hexaploid F_2_ mapping populations LS783 x YS783 (population size: 193) and LS665 x LS783 (population size: 182) to evaluate the homologous recombination frequency (RF) of AB and D genomes. Meanwhile, the RF of the other two F_2_ populations of *Ae. tauschii* SQ665 x SQ783 (population size: 124) and *T. turgidum* Langdon x Yuanwang (population size: 192) were calculated as a control to investigate the RF changes of the diploid and tetraploid genomes after their hexaploidization, respectively.

### SNP genotyping

Among the four F_2_ genetic populations, the two populations SQ665 x SQ783 (D_1_D_1_ x D_2_D_2_) and LS665 x LS783 (AABBD_1_D_1_ x AABBD_2_D_2_) were grouped to the first genetic population set so as to analyze the RF change of the ancestral diploid genome after its hexaploidization. The other two F_2_ populations Langdon x Yuanwang (A_1_A_1_B_1_B_1_ x A_2_A_2_B_2_B_2_) and LS783 x YS783 (A_1_A_1_B_1_B_1_DD x A_2_A_2_B_2_B_2_DD) was set together for the RF analysis of the ancestral tetraploid genome after their hexaploidization. And the 4 populations and their involved parents were planted in the field using the single-seed precision sowing by hand. The distance between the rows was 30 centimeter and the distance between seeds in each row was 20 cm. 50 mg of plant tissue was collected from 2-week-old seedlings and their DNA was extracted using the NuClean Plant Genomic DNA Kit (CWBio, Beijing, China). Eluted DNA was quantified using Qubit 4 Fluorometer (Life Technologies Holdings Pte Ltd, Singapore) and then normalized using a 12-channel electronic pipette with a volume range of 10 to 100µL (Eppendorf, Hamburg, Germany) to obtain the concentration required for genotyping.

For the first genetic population set and their parents Langdon, SQ665, SQ783 and two SHWs LS665 and LS783, their genotyping by sequencing (GBS) analysis was performed by using DArT-Seq™ technology relying on a complexity reduction method to enrich genomic representations with single copy sequences and subsequently performing next-generation sequencing using HiSeq2500 (Illumina, USA), for obtaining sequencing reads and detecting SNPs by comparing to the RefSeq v1.0 assembly of CS (IWGSC et al., 2018). About 100 μl of 50 ng μl^-1^ DNA sample was sent to Diversity Arrays Technology P/L (Bruce, Australia) for SNP and DArT analysis. However, only SNP data was used for linkage map construction and RF calculation in this study, as co-dominant molecular data had advantage on RF calculation that was more accurate than dominant molecular data (Sun et al., 2012). Among the 28331 detected SNPs after comparing obtained sequence reads to CS assembly (IWGSC et al., 2018), a total of 9043 SNP labels were evenly discovered and theoretically distributed throughout the diploid D genome. In order to confirm their site specificity on D genome, the *durum* wheat Langdon as the same AABB donor of both LS665 and LS783 was used to delete these SNPs also detected in the AB genome of Langdon out from the 9043 SNPs on the diploid D genome.

For the second genetic population set and all their parents including SQ783, Langdon, Yuanwang, LS783 and YS783, their genotyping were executed on the Affymetrix platform of Axiom Wheat Breeder’s Genotyping Array with 13947 SNP markers developed by China Golden Marker Biotech Co Ltd (Beijing, China). The collected fluorescence signal from SNP array were processing and analyzed by the functions of apt-genotype-axiom for genotype calling, ps-metrics for generating various QC metrics and ps-classification for classifying SNPs in the software of Affymetrix Axiom Analysis Suite version 4.0.1. Among the 13947 SNP markers, a total of 9487 SNP labels fixed on the chip is designed to be distributed on AB genomes, and SQ783 was serviced to remove these SNP markers with their fluorescence signal also detected on the D genome of *Ae. tauschii*.

### Genetic maps and recombination frequency

For the D genome of the first genetic population set, a total of 744 polymorphic SNPs were remained for linkage map construction and RF calculation, and the defined rules filtering the detected SNPs generally applied to further analysis were as follows: (1) the best Blastn hits with top score was unique in the D genome when using allele reference sequences of SNP assay querying the *A. tauschii* AL8/78 genome sequence assembled by Luo et al. (2017); (2) the remained SNP markers must have no genotype (displaying as missing data) on the AB genome of Langdon; (3) For a employed SNP site, their polymorphism should also be detected between the parents of both diploid and hexalpoid F_2_ populations; (4) genotypes of all involved parents was homozygous at the workable SNP sites, and the diploid parents SQ665 and SQ783 had the same genotypes with their SHWs LS665 and LS783, respectively; (5) the SNP markers could detected both homozygous and heterozygous genotypes in both populations. The reserved SNPs must obey all above-mentioned rules simultaneously. For the AB genome, a total of 904 SNP markers were selected for linkage map construction and RF calculation, obeying the basic principles of the above-mentioned rules in filtering process of SNPs using the CS IWGSC RefSeq v1.0 assembly (IWGSC et al., 2018) and their own involved parents.

The QTL IciMapping Software version 3.2 (Li et al., 2007; Meng et al., 2015) was used for genetic linkage map construction and calculation of the RF between adjacent SNP. The maximum likelihood method was used to estimate RF by the QTL IciMapping Software version 3.2 (Sun et al., 2012). In order to comparing the RF between two genetic populations of each set, the SNP markers for genetic map construction were aligned using their physical positions on the CS RefSeq v1.0 assembly (IWGSC et al., 2018) for AB genome and sequence assembly of *Ae. tauschii* AL8/78 (Luo et al., 2017) for D genome, and then the RF between two adjacent SNP markers in physical map was calculated using the functional of ‘algorithm by input’ in the software.

## Results

### Distribution of polymorphic SNPs on AB and D genomes

Wheat Breeder’s Genotyping Array and DArT-Seq™ technology were used to genotype the AB and D genomes of two genetic population sets and their parents in this study, respectively. For AB genome’s genotyping, a total of 9925 designed SNP labels anchored on Wheat Breeder’s Genotyping Array, and their numbers ranged from 463 of 6A to 997 of 3B distributed on chromosomes of AB genome (Table 1). And about 10-30 SNPs per 20 Mb were distributed on AB genome, and the pericentromeric SNPs was less than other genomic regions, especially on 1B, 2B, 4B and 5A chromosomes (Fig. 1A). For D genome, 9043 SNPs was totally obtained by DArT-Seq™ technology, based on genotyping by sequencing (GBS) and aligning the Illumina sequencing reads to the RefSeq v1.0 assembly of CS. The SNP numbers distributed on chomosomes of D genome ranged from 795 on 4D to 1850 on 2D (Table 1). Their distributions were quietly disequilibrium along the chromosomes of D genome. The amounts of detected SNPs increases consistently and linearly with the physical distance away from the centromere of each chromosome, and obtained SNP sites were much more in distal centromere-free regions (Fig. 1B).

**Fig. 1.**
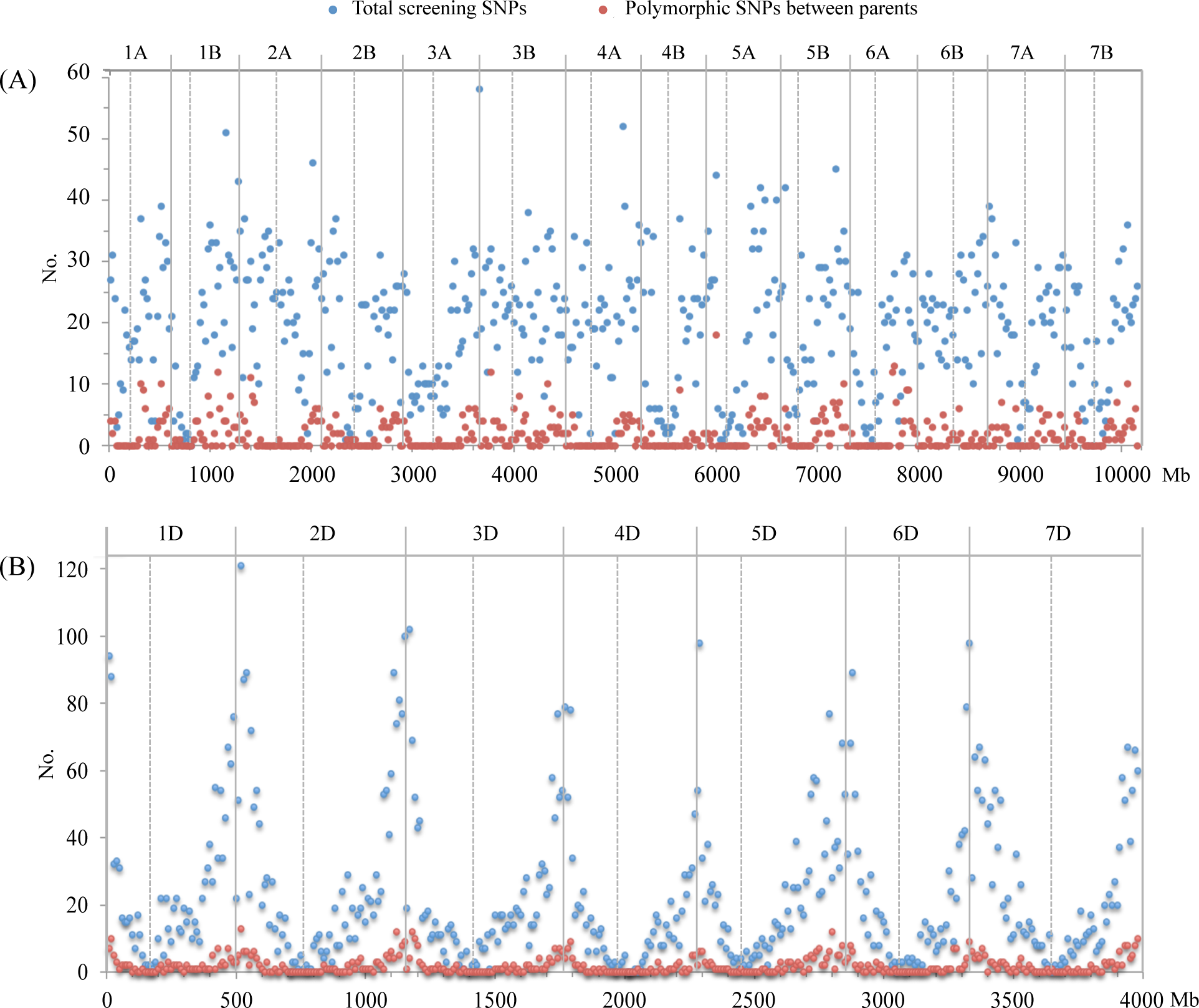
Amounts of the screening SNP markers used for genotyping and polymorphic SNPs number between the parents. (A) AB genome with statistical window sizes of 20 Mb; (B) D genome with statistical window sizes of 10 Mb. The centromere of each chromosome is marked by dotted line.

**Table 1.**
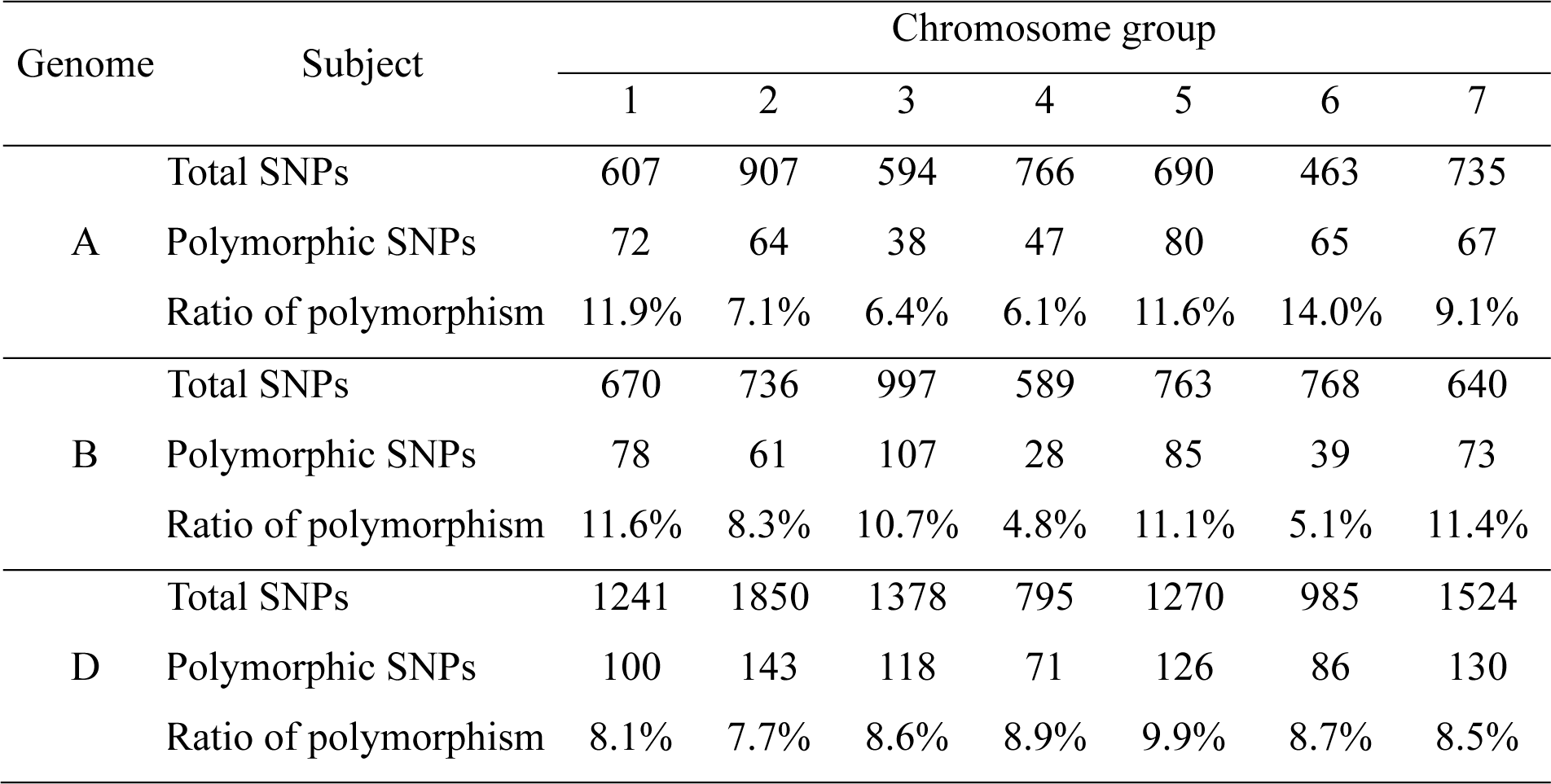
Distribution of SNP markers in A, B and D genome and their polymorphism between genotyped parents.

Obeying the polymorphic SNP filtering rules mentioned in M&M, a total of 774 polymorphic SNPs were detected in D genome of the first genetic population set with their parental dipliody and SHWs, the numbers of polymorphic SNPs on the chromosomes ranged from 71 of 4D to 143 of 2D (Table 1). Distributions of polymorphic SNPs on chromosomes showed a linear decrease when approaching the centromeres (Fig. 1B). The polymorphic SNPs around centromere was nearly blank in the first genetic population set, and the majority of the polymorphic SNPs were clustered in the genomic regions away from the pericentromeric regions and covered about 1,520 Mb totally with at least 2 SNPs per 10 Mb on average (Fig. 2).

**Fig. 2.**
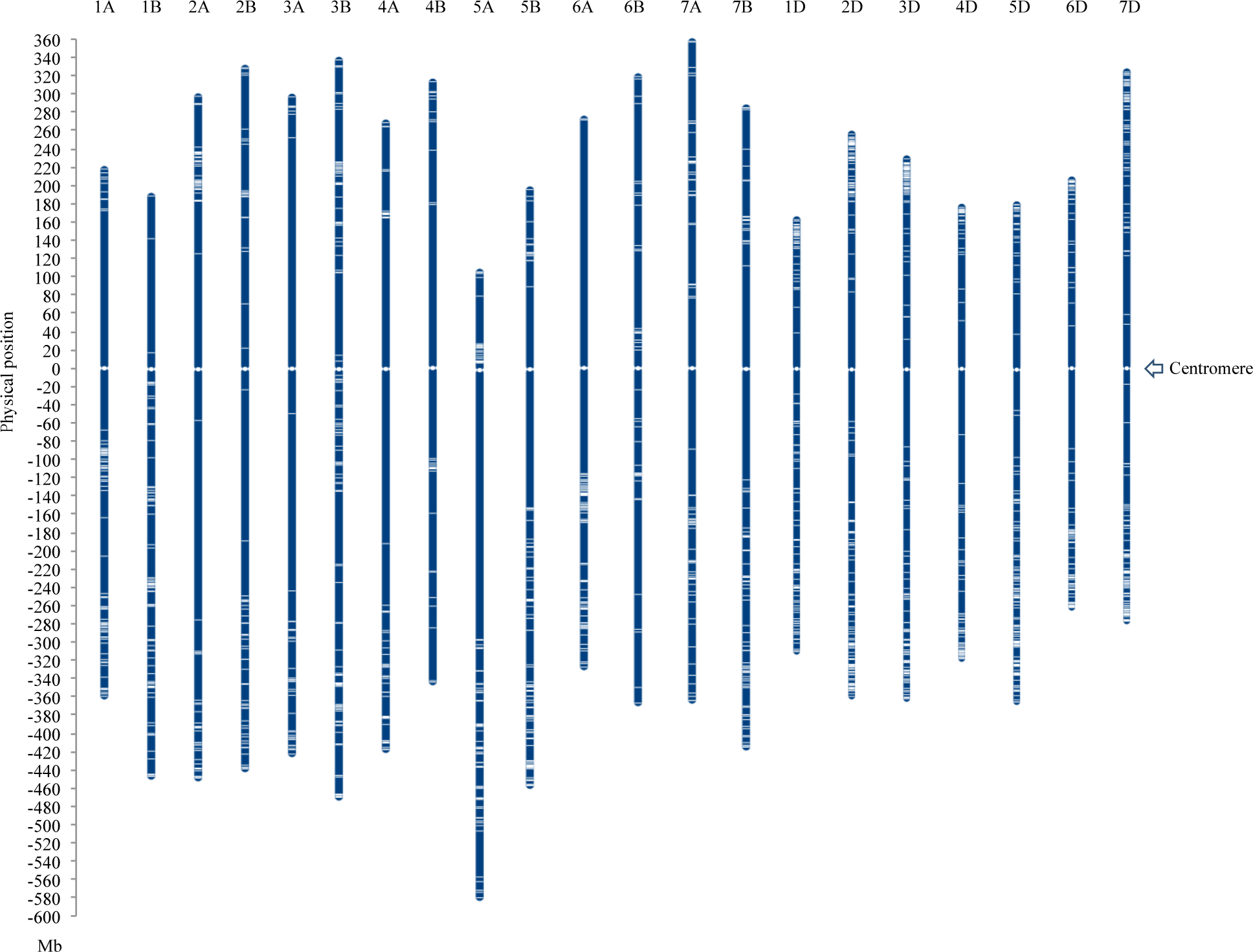
Distribution of the polymorphic SNPs on the physical maps of AB and D genome. The physical position of the centromere on each chromosome was marked as 0 Mb.

For AB genome, 9487 SNPs on chip were selected artificially by China Golden Marker Biotech Co Ltd and distributed more equilibriously in AB genome than D genome (Fig. 1). And a total of 904 polymorphic SNPs were identified between two tetraploid parents and their SHWs, and the number of polymorphic SNPs detected on each chromosome ranged from 28 on 4B to 107 on 2B (Table 1). The polymorphic SNPs was also very rare in the centromere region of all chromosomes except chromosome 3B possessing more polymorphic SNPs in pericentromeric regions. Moreover, the polymorphic SNPs were detected to be clustering in some genomic regions of AB genome, i.e., the 80-140 Mb region on the long arm of chromosome 1A, the 190-210 Mb region on the short arm of chromosome 2A (Fig. 1A, Fig. 2). The genomic regions clustering the majority polymorphic SNPs covered total physical genome length of about 2,320 Mb with at least 4 SNPs per 20 Mb on average (Fig. 2).

### The level of genetic recombination among tetraploid genome after hexaploidization

For tetraploid wheats (AABB) and their SHWs, all polymorphic SNPs covered total physical lengths of 4771.2 Mb and 4944.2 Mb on A and B genomes, respectively. And the physical lengths covered by polymorphic SNP markers on each chromosome ranged from 581.5 Mb of 1A to 813.7 Mb of 3B (Table 2). The genetic maps for both tetraploid and hexaploid populations were constructed based on the physical positions of the 904 polymorphic SNPs. For the tetraploid population, the total genetic map length of AB genome was 3135.9 cM, and the length of each chromosome ranged from 131 cM of 1B to 537 cM of 4B (Fig. 3, Table 4). For their SHW-derived population, the total genetic map length of AB genome was 2326.7cM, and the length of each chromosome ranged from 109.5cM of 1B to 394.5cM (Fig. 3, Table 4). The map lengths of chromosome 2B (393.0 cM) and 4B (537.9 cM) calculated using tetraploid population was 112.7 cM and 143.3 cM longer than those (280.3 cM for 2B, 394.5 cM for 4B) calculated in their SHW-derived population, respectively (Fig. 3, Table 4). Obviously, the length difference of 2B and 4B genetic maps between the tetraploid and hexaploid populations was mostly caused by linkage gaps existing on their maps. Furthermore, the linkage gaps on chromosome 4B map constructed using the tetraploid population were more than that constructed using the haxaploid population (Fig. 3, Table 4).

**Fig. 3.**
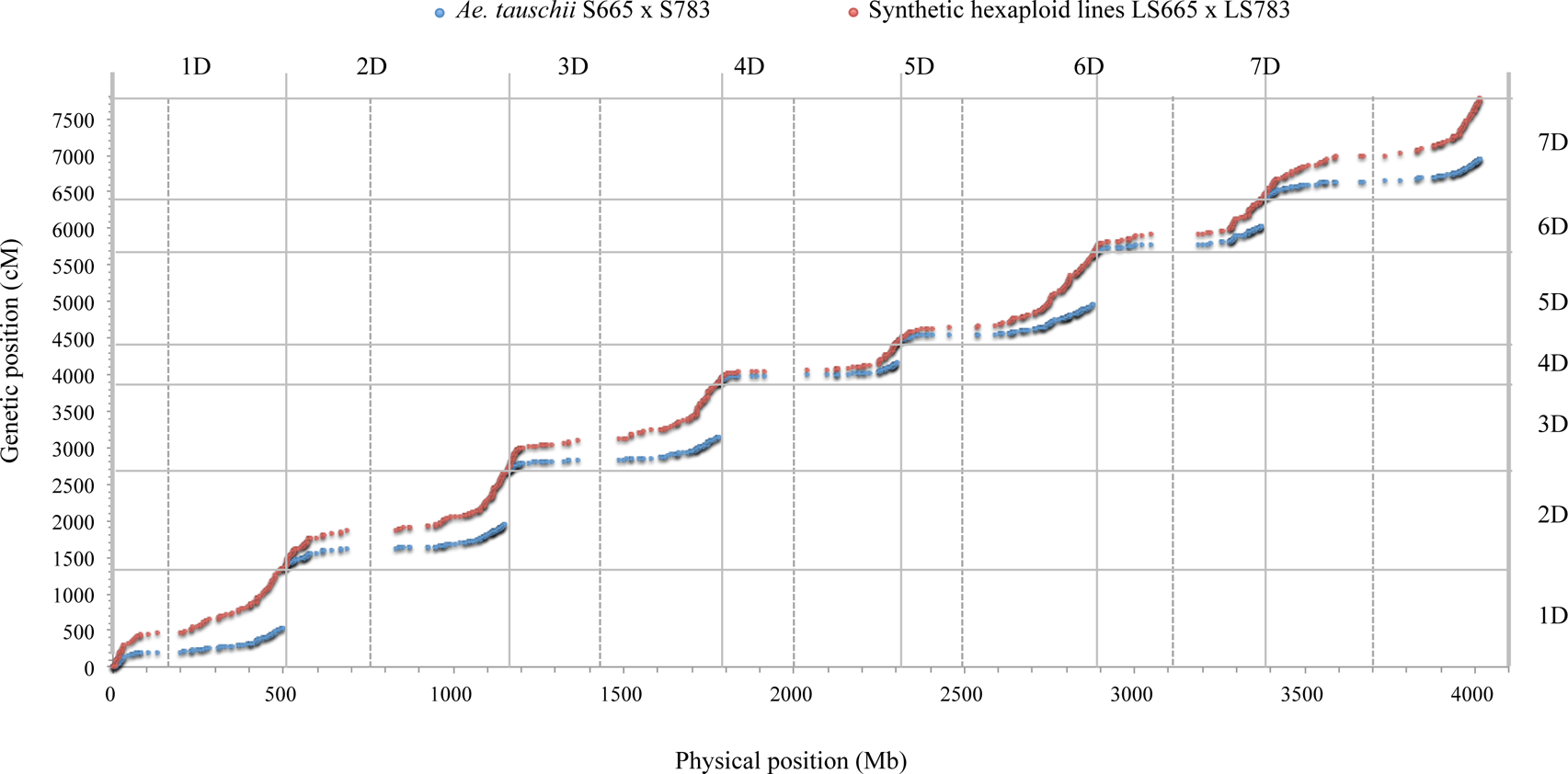
The collinearity between genetic position and physical position of polymorphic SNPs in both diploid *Ae. tauschii* and their SHW-derived F_2_ populations. The centromere of each chromosome is marked by dotted line.

**Table 2.** Genetic interval length and RF between two adjacent and linked SNP loci on each chromosome of AB genome in tetraploid wheat and their SHW F_2_ populations

| Genome | Group | SNPs No. | Covering length<br>(Mb) | Genetic interval length (cM) |  |  |  | Recombination frequency |  |  |  |
| --- | --- | --- | --- | --- | --- | --- | --- | --- | --- | --- | --- |
| | | | | TTP | SHWP | $\Delta_{\text{SHW-TT}}$ | P value | TTP | SHWP | $\Delta_{\text{SHW-TT}}$ | P value |
| A | 1 | 72 | 581.5 | 2.23 | 2.03 | -0.20 | 0.77 | 2.18% | 2.00% | -0.18% | 0.78 |
|  | 2 | 64 | 751.5 | 2.61 | 1.98 | -0.63 | 0.45 | 2.54% | 1.94% | -0.60% | 0.46 |
|  | 3 | 38 | 725.3 | 4.33 | 2.92 | -1.41 | 0.34 | 4.10% | 2.88% | -1.21% | 0.36 |
|  | 4 | 47 | 691.4 | 2.88 | 2.08 | -0.80 | 0.32 | 2.82% | 2.07% | -0.75% | 0.34 |
|  | 5 | 80 | 687.7 | 2.49 | 1.54 | -0.95 | 0.21 | 2.38% | 1.51% | -0.87% | 0.21 |
|  | 6 | 65 | 605.3 | 1.13 | 1.14 | 0.01 | 0.99 | 1.11% | 1.12% | 0.01% | 0.98 |
|  | 7 | 67 | 728.5 | 2.90 | 2.48 | -0.41 | 0.72 | 2.70% | 2.36% | -0.34% | 0.74 |
| B | 1 | 78 | 639.2 | 1.70 | 1.42 | -0.28 | 0.64 | 1.66% | 1.40% | -0.26% | 0.65 |
|  | 2 | 61 | 773.7 | 1.83 | 1.31 | -0.52 | 0.29 | 1.81% | 1.31% | -0.50% | 0.29 |
|  | 3 | 107 | 813.7 | 1.36 | 1.47 | 0.10 | 0.86 | 1.28% | 1.43% | 0.15% | 0.76 |
|  | 4 | 28 | 663.5 | 2.29 | 2.78 | 0.49 | 0.72 | 2.23% | 2.74% | 0.50% | 0.70 |
|  | 5 | 85 | 656.4 | 1.88 | 1.51 | -0.36 | 0.47 | 1.84% | 1.50% | -0.34% | 0.49 |
|  | 6 | 39 | 692.3 | 2.74 | 1.45 | -1.29 | 0.22 | 2.64% | 1.44% | -1.20% | 0.23 |
|  | 7 | 73 | 705.4 | 2.05 | 1.17 | -0.88 | 0.22 | 1.93% | 1.16% | -0.77% | 0.22 |

For AB genome, the average genetic distance between two adjacent and linked SNP loci in different chromosomes ranged from 1.13 cM of chromosome 6A to 4.33 cM of chromosome 3A in the tetraploid F_2_ population, and from 1.14 cM of 6A to 2.92 of 3A in their hexaploidy population (Table 2). The average genetic distance between two adjacent and linked SNP loci in tetraploid genome increased by 0.01 cM on chromosome 6A, 0.10 cM on chromosome 3B and 0.49 cM on chromosome 6A after hexaploidization, while the average genetic distance decreased by 0.20 cM to 1.41 cM in other chromosomes. However, no significant difference on the average genetic distance between two adjacent and linked SNP loci on each chromosome have been found between tetraploid wheat and their hexaploidy (Table 2). The average RF between two adjacent and linked SNP loci on different chromosomes had also no significant difference between tetraploid wheat and their hexaploidy (Table 2), and their Δ_SHW-TT_ of RF were mostly distributed from −0.05% to 0.03% on total AB genome (Fig. 5), suggesting that no remarkable change of RF had happened in most physical region of tetraploid genome after their hexaploidization.

**Fig. 4.**
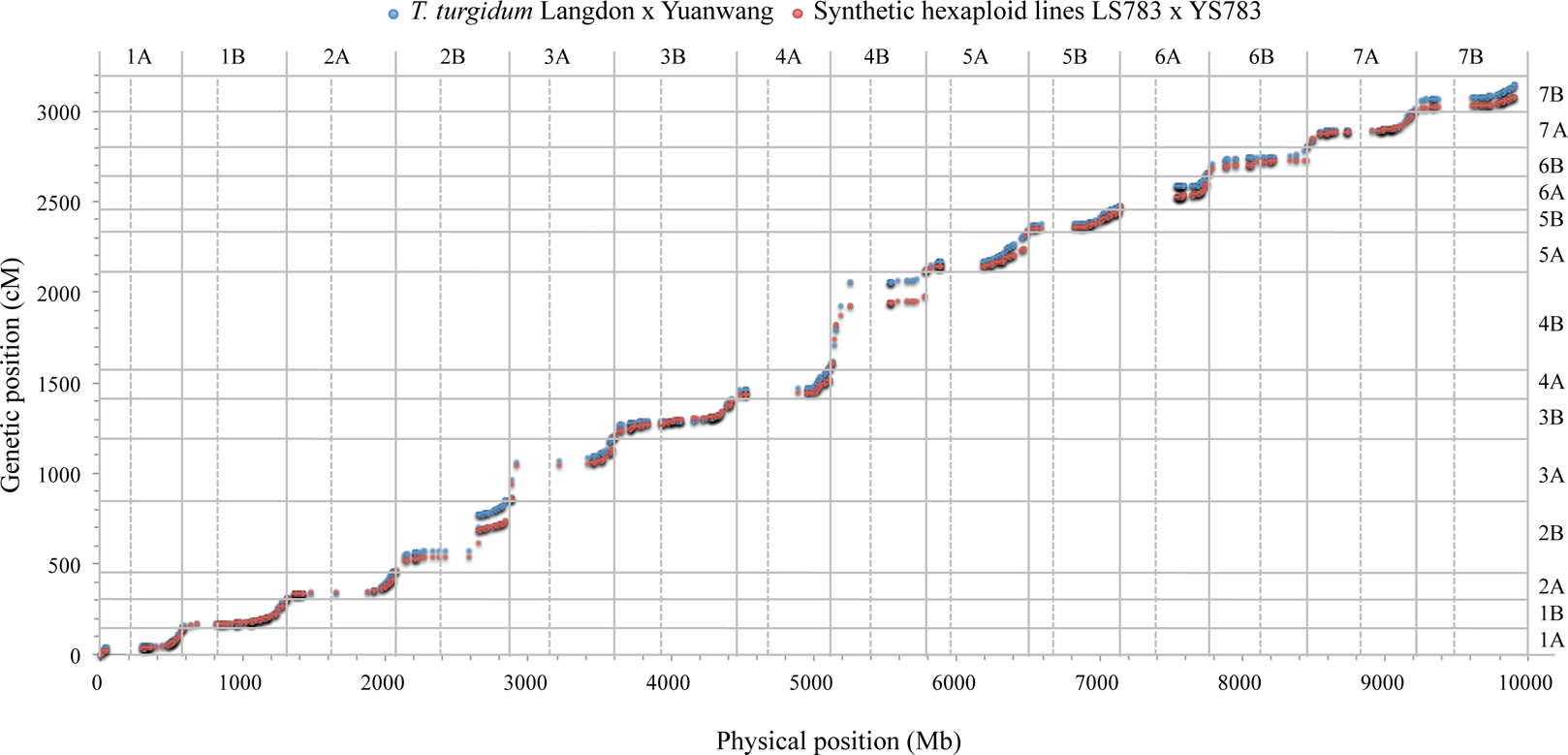
The collinearity between genetic position and physical position of polymorphic SNPs in both tetraploid wheat and their SHW-derived F_2_ populations. The centromere of each chromosome is marked by dotted line.

**Fig. 5.**
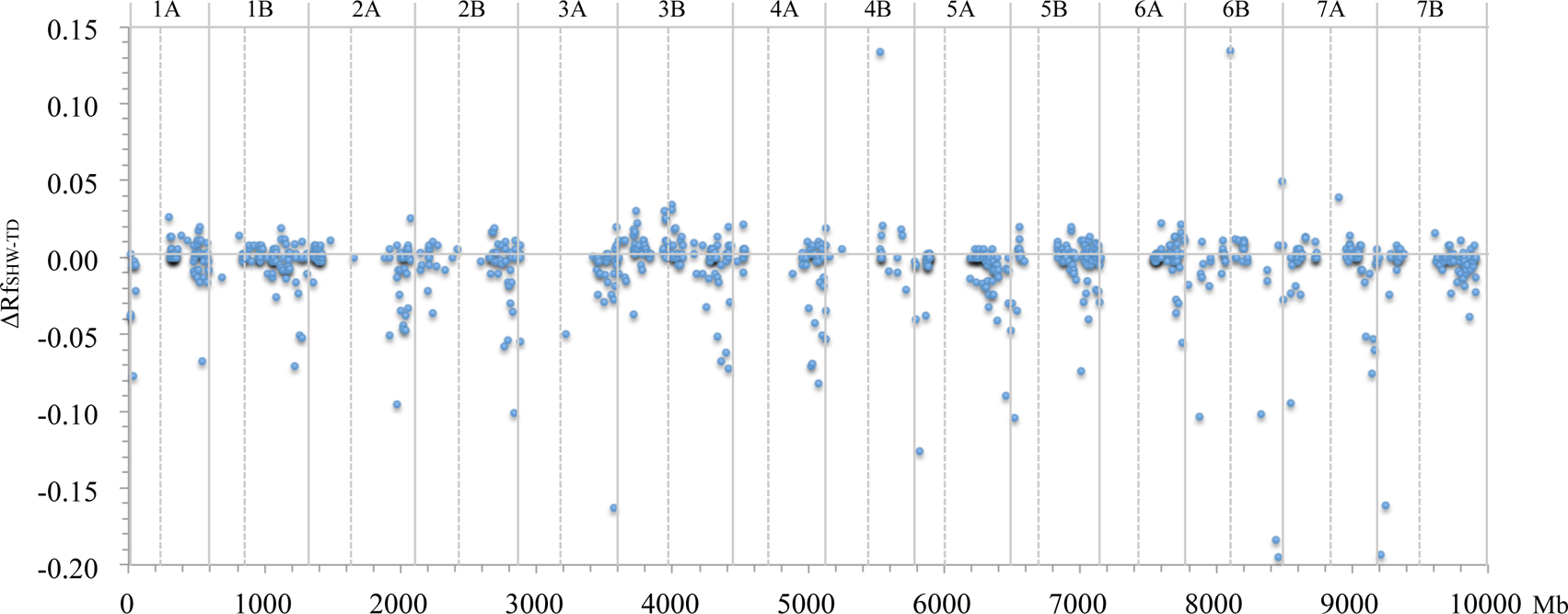
The RF change between tetraploid wheat (AABB) and their SHW-derived F_2_ populations. The horizontal axis value indicate the physical position of the right boundary of the interval between two adjacent and linked SNP loci.

However, some genomic regions of tetraploid genome (AB) were also found to have larger RF changes after hexaploidization (Fig. 5). In the pericentromeric region of 4B, the RF between the physical position of 137.6 Mb and the physical position of 414.7 Mb increased by 13.4% from 1.5% in tetraploid wheat to 14.9% in their SHW. And in the centromere region of 6B, the RF was also increased by 13.4% under the hexaploid genetic background (Fig. 5), suggesting that hexaploidization may increase the RF in pericentromeres of specific chromosomes. Newtheless, the RF between the physical position of 705.3Mb and 729.4Mb on chromosome 3A of tetraploid wheat decreased by 16.3% when introduced into a hexaploid genetic background. And the RFs of the genomic regions closed to telomere of chromosome 5A and 5B decreased by more than 10% after hexaploidization. The RFs of the intervals from 617.8Mb to 696.9Mb on long arm of chromosome 6B and from 19.3Mb to 65.1Mb on short arm of chromosome 7B decreased by more than 15% in a hexaploid genetic background (Fig. 5).

### The level of genetic recombination among diploid genome after hexaploidization

The polymorphic SNPs totally covered about 3992.9 Mb of physical map of *Ae. tauschii* and ranged from 493.8 Mb of 6D to 648.7 Mb of 2D on chromosomes (Table 3). The genetic map constructed by the diploid population was totally 3334.9 cM long, and the map length of each chromosome ranged from 263.2 cM of 4D to 593.3 cM of 2D (Table 5, Fig. 4). The total length of genetic map constructed by the hexaploidy population was 7185.4 cM, two times more than the genetic map of *Ae. Tauschii*, which also happened on each chromosome map (Fig. 4). And any linkage gaps did not exit on both chromosome maps of D genome constructed by *Ae. tauschii* and their hexaploid populations (Table 5, Fig. 4).

**Table 3.** Genetic interval length and RF between two adjacent and linked SNP loci on each chromosome of D genome in diploid *Ae. Tauschii* (ATP) and their SHW (SHWP) F_2_ populations

| Chromosome | SNPs No. | Covering length | Genetic interval length (cM) |  |  |  | Recombination frequency |  |  |  |
| --- | --- | --- | --- | --- | --- | --- | --- | --- | --- | --- |
| | | (Mb) | ATP | SHWP | $\Delta_{SHW-AT}$ | P value | ATP | SHWP | $\Delta_{SHW-AT}$ | P value |
| 1D | 100 | 497.9 | 5.39 | 13.67 | 8.28 | 0.00 | 5.34% | 13.17% | 7.83% | 0.00 |
| 2D | 143 | 648.7 | 4.18 | 9.20 | 5.02 | 0.00 | 4.14% | 9.00% | 4.86% | 0.00 |
| 3D | 118 | 623.0 | 4.17 | 10.56 | 6.39 | 0.00 | 4.14% | 10.28% | 6.14% | 0.00 |
| 4D | 71 | 520.9 | 3.76 | 7.83 | 4.07 | 0.00 | 3.74% | 7.70% | 3.96% | 0.00 |
| 5D | 126 | 572.6 | 4.20 | 9.56 | 5.36 | 0.00 | 4.16% | 9.37% | 5.21% | 0.00 |
| 6D | 86 | 493.8 | 4.57 | 9.04 | 4.47 | 0.00 | 4.53% | 8.87% | 4.33% | 0.00 |
| 7D | 130 | 636.0 | 4.21 | 10.57 | 6.36 | 0.00 | 4.18% | 10.26% | 6.08% | 0.00 |

**Table 4.** Genetic lengths of chromosome maps aligned with physical map for AB genome and self-organized by nnTwoOpt method.

| Chromosome | Marker No. | TDP (cM) |  | SHWP (cM) |  |
| --- | --- | --- | --- | --- | --- |
|  |  | Self-organization | Physical alignment | Self-organization | Physical alignment |
| 1A | 72 | 157.5 | 158.1 | 134.0 | 144.2 |
| 1B | 78 | 130.7 | 131.1 | 104.0 | 109.5 |
| 2A | 64 | 151.5 | 164.7 | 109.2 | 124.9 |
| 2B | 61 | <u>298.5 (2<sup>a</sup>)</u> | <u>393.0 (3)</u> | <u>196.2 (2)</u> | <u>280.3 (3)</u> |
| 3A | 38 | <u>262.6 (1)</u> | <u>348.7 (2)</u> | <u>220.6 (1)</u> | <u>281.2 (2)</u> |
| 3B | 107 | <u>202.1 (1)</u> | <u>202.8 (1)</u> | 152.7 | 181.1 |
| 4A | 47 | 172.0 | <u>179.6 (1)</u> | 109.7 | 115.4 |
| 4B | 28 | <u>299.1 (3)</u> | <u>537.9 (5)</u> | <u>244.6 (2)</u> | <u>394.5 (3)</u> |
| 5A | 80 | 195.4 | 196.4 | 119.8 | 121.7 |
| 5B | 85 | 157.1 | 157.7 | 120.5 | 127.1 |
| 6A | 65 | <u>162.6 (1)</u> | <u>180.6 (1)</u> | <u>118.4 (1)</u> | <u>123.9 (1)</u> |
| 6B | 39 | 146.6 | 146.9 | 65.4 | 74.5 |
| 7A | 67 | 190.8 | 191.1 | 155.0 | 163.9 |
| 7B | 73 | 126.4 | 147.4 | 81.3 | 84.4 |
| Total | 904 | 2653.0 | 3135.9 | 1931.5 | 2326.7 |

<sup>a</sup>Number of map gaps with genetic distance more than 50cM.

**Table 5.** Genetic map lengths for chromosome maps of D genome aligned with physical map and self-organized by nnTwoOpt method.

| Chromosome | Marker No. | ATP (cM) |  | SHWP (cM) |  |
| --- | --- | --- | --- | --- | --- |
|  |  | Self-organization | Physical alignment | Self-organization | Physical alignment |
| 1D | 100 | 520.6 | 533.6 | 1148.8 | 1227.3 |
| 2D | 143 | 548.8 | 593.3 | 1157.1 | 1194.6 |
| 3D | 118 | <u>506.7 (1<sup>a</sup>)</u> | 488.4 | 1116.2 | 1135.3 |
| 4D | 71 | 251.4 | 263.2 | 515.0 | 519.8 |
| 5D | 126 | 504.1 | 524.5 | 1074.8 | 1105.3 |
| 6D | 86 | 368.0 | 388.6 | 706.2 | 723.6 |
| 7D | 130 | 527.1 | 543.3 | 1260.6 | 1279.4 |
| Total | 774 | 3226.7 | 3334.9 | 6978.7 | 7185.4 |

<sup>a</sup>Number of linkage gaps with genetic distance more than 50cM.

For the diploid *A. tauschii* population, the average genetic lengths between two adjacent SNPloci on each chromosome ranged from 3.76 cM of 4D to 5.39 cM of 1D, while it ranged from 7.83 cM of 4D to 13.67 cM of 1D in their SHW-derived population (Table 3). For each chromosome, the average genetic length between two adjacent SNP loci on the hexaploid genetic background was significantly over 2-fold longer than that in the diploid, and their differential value Δ_SHW-AT_ between the diploid and hexalpoid populations ranged from 4.07 cM on 4D to 8.28 cM on 1D (Table 3). The average RF between two adjacent SNP loci in the hexaploid population was also significantly over 2-fold higher than that in the diploid population (Table 3). Their average Δ_SHW-AT_ of RF on each chromosome between two populations with different ploidy levels ranged from 3.96% of 4D to 7.83% of 1D (Table 3). The Δ_SHW-AT_ of RF between two adjacent SNP loci was mostly distributed from −5% to 17% along the whole D genome (Fig. 6), with average mean of 5.5% and ratio of 2.3 (Table 3: RF_SHWP_/RF_ATP_). These results suggested that the hexaploidization enhancing RF of the ancestral diploid genome.

**Fig. 6.**
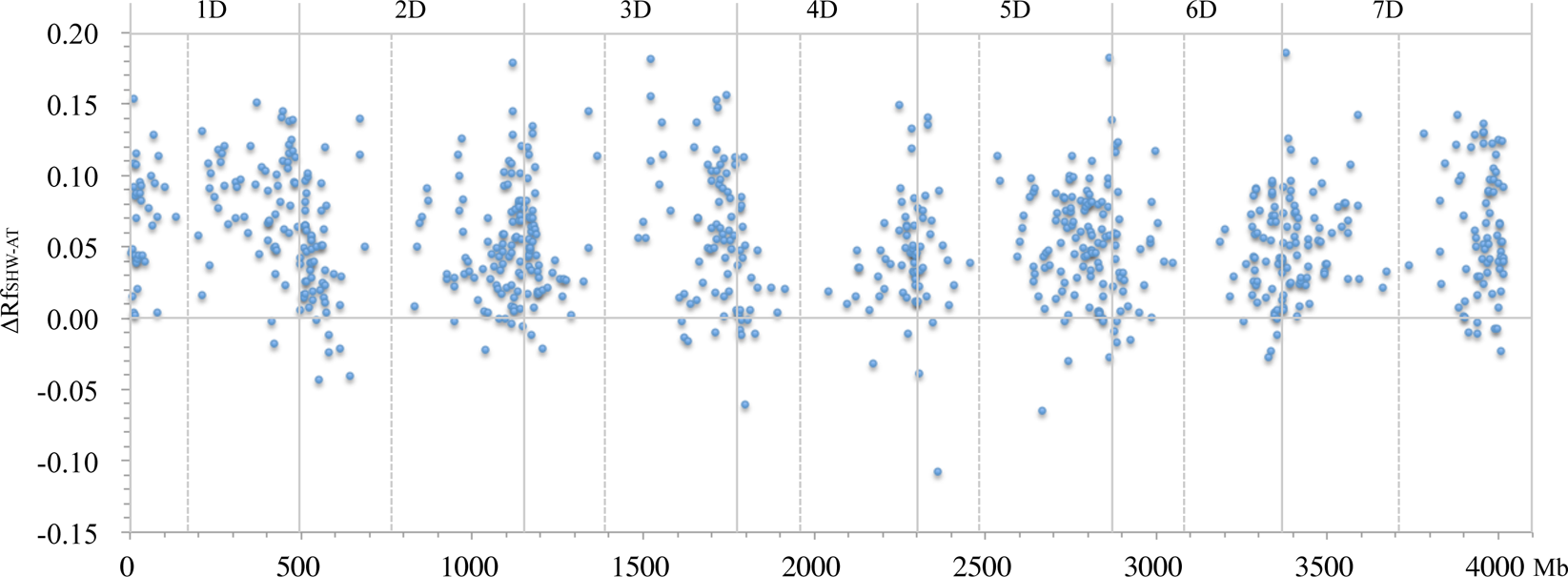
The RF change between diploid *Ae. tauschii* and their SHW-derived F_2_ populations. The horizontal axis value indicate the physical position of the right boundary of the interval between two adjacent and linked SNP loci.

In addition, 7 outliers were also observed on some genomic regions, with a stem-and-leaf plot displaying the distribution of Δ_SHW-TD_ of RF. On the region from the physical position 616.76 Mb to 618.10 Mb on chromosome 2DL, the RF between two adjacent SNP loci increased by 17.94 % after their hexaploidization (Fig. 6). On the physical regions of 369.26 Mb. - 371.13 Mb on 3D, the intervals obtained increased RFs by 18.21% after hexaploidization. And the intervals of 561.63 Mb - 562.21 Mb on chromosome 5D and 5.34 Mb - 7.21 Mb on 7D obtained increased RFs by 18.23% and 18.57% after hexaploidization, respectively (Fig. 6). However, 3 genomic regions’ RFs decreased by more than 6% after hexaploidization. In the interval from the physical position of 45.65 Mb to 58.27 Mb on chromosome 5DS, the RF decreased by 10.74% after hexaploidization, and the intervals of 17.58 Mb - 20.81 Mb on 4D and 352.32 Mb - 364.72 Mb on 5D obtained 6.03% and 6.52%-decreasing RFs after hexaploidization, respectively (Fig. 6).

## Discussion

### The genetic maps constructed based on physical positions of SNPs

In this study, SNPs used for constructing the genetic maps of AB genome and D genome were aligned based on their physical positions on the sequence assembly of Chinese Spring (IWGSC et al., 2018) and *A. tauschii* AL8/78 (Luo et al., 2017), respectively. Moreover, for genetic map construction, nearest neighbor and two-opt (nnTwoOpt) were also used for tour construction and its improvement, which is similar to Travelling Salesman Problem (TSP) (Lin and Kernighan, 1973). For AB genome, chromosomes without any linkage gaps, such as 1A, 1B, 2A, 5A, 5B, 6B, 7A, and 7B, almost had the same genetic lengths of chromosome maps self-organized by nnTwoOpt method to those aligned with physical map CS (Table 4). For the chromosomes 2B, 3A, 3B, 4B and 6A, linkage gaps were found in these chromosomes and caused by the extreme disequilibrium of SNP-polymorphism distribution. For D genome, the polymorphism SNPs on each chromosome were much more than that in AB genome, and only one linkage gap on 3D was found on the chromosome maps self-organized by nnTwoOpt method among the whole genome (Table 5). And only small difference was found in genetic lengths between self-organizing genetic maps and that aligned with physical map (Table 5). So, we fixed the order of SNPs aligned with their physical position in CS for AB genome and AL8/78 for D genome, in order to investigate the change of the genetic length and RF of the diploid and tetraploid genome after their hexaploization directly and conveniently.

More over, linkage gaps were found on some chromosomes of the tetraploid genome, and their genetic distance between two adjacent SNP loci was more than 50 cM (Table 4). More double-crossovers exist between two adjacent SNP loci in the physical map, if their genetic distance was more than 50cM. And the measured RF could not reflect the true recombination frequency, which might be much more larger than the calculated value. Considering this, we only compared the genetic distance and RF between two SNP loci that were both adjacent and linked in the maps.

### Enhanced genetic recombination of ancestral diploid genome after hexaploidization

In chromosome level, we compared the average RF betweentwo adjacent and linkage SNP loci in diploid and tetraploid populations with their SHW-derived population. And significant increase of RF was observed only in the diploid D genome after their hexaploidization, but not in the tetraploid AB genome. These results suggested that the extent of RF increase in ancestral genome depended on their level of polyploidization, while the increase rate of RF from diploidy to hexaploidy was much more than that from tetraploidy to hexaploidy. In *Brassica rapa* L., the total genetic length of A7 linkage group increased from 52 cM to 96 cM after their tetraploidization from diploidy to tetraploidy (Leflon et al., 2010). In *Arabidopsis thaliana*, the meiotic recombination frequency increased from 15.4% of diploid *A. thaliana* to 24.1% of allotetraploid *A. suecica* or 20.5% of autotetraploid *A. thaliana* (Pecinka et al., 2011). These reported data showed that the increase rate of RF was about 100% from diploidy to tetraploidy, and the RF increased more than 200% from diploidy to hexaploidy (Table 3).

However, this situation might be only suitable for euploidy but not for aneuploidy, and in *B. rapa*, A7 linkage group of allotriploid got 4-fold increase of the total genetic length more than both diploid and tetraploid (Leflon et al., 2010). Aneuploidy causes greater genome instability than polyploidy for organisms (Birchler et al., 2001; Chen, 2007) and aneuploidy itself can be responsible for the procreation of chromosomal instability (Potapova et al., 2013; Dürrbaum and Storchová, 2016). Chromosomes that remain as univalents in the aneuploidy could lead to a compensatory increase in crossover frequency among unaffected bivalents in aneuploidy (Parker, 1975; Tease and Jones, 1975; Carlton et al., 2006; Leflon et al., 2010).

The RF of the diploid D genome was significantly enhanced after their hexaploidization, and the genetic mechanisms for their increase has not been clear yet. QTLs for crossover (CO) has been detected in hexaploid wheat, and most detected QTL were distributed on AB genome. With 13 recombinant inbred mapping populations, Gardiner et al. (2019) detected 5 QTLs for CO frequency on 2A, 2B, 4B, 5A and 6A, respectively; Jordan et al. (2018) detected 40 QTLs for total CO frequency by nested association mapping, and most of them were mapped to AB genome. These results tended to make an indication that the genetic factors might exist in AB genome, which increased the RF of diploid D genome after their hexaploidization with AB genome. However, the phenotypic effect of these detected QTLs in increasing the total CO frequency were < 10.5% that was much less than the increase extent of >200% for RF caused by hexaploidization, suggesting that the increase of RF in our study was mostly caused by hexaploidization much more than the genetic factors existed on AB genome.

### Polyploidization enhancing variation and adaptive evolution of bread wheat

Allopolyploidy induced rapid revolution to wheat often by two ways: (1) allopolyploidization triggers rapid genome changes through the instantaneous generation of a variety of cardinal genetic and epigenetic alterations, which can generate heterosis in polypoid plants (Chen and Ni, 2006; Chen, 2007), and (2) the allopolyploid condition facilitates sporadic genomic changes during the life of the species that are not attainable at the diploid level, which is caused by gene redundancy (Feldman et al., 1997; Ozkan et al., 2001; Shaked et al., 2001; Feldman and Levy, 2005; Comai, 2005). *Ae. tauschii*, the D genome donor of the common wheat, was regarded as a gene repertoire for wheat adaptation. By hexaploidization, *Ae. tauschii* had been added in to the genome of tetraploid wheat, and made the hexaploid wheat being a major type of cultivated wheat, which accounting for about 95% of world wheat production, while the tetraploid wheat only accounting for the other 5% (Dubcovsky and Dvorak, 2007), suggesting that the adding of D genome of *Ae. tauschii* made bread wheat more adaptive to alterable environments and then spread more rapidly than the tetraploid wheat.

*Ae. tauschii* is a widely distributed (van Slageren, 1994) and genetically diverse species (Lubbers et al., 1991; Dvorak et al., 1998; Schneider et al., 2008), with superiority in resistance to disease, pest and tolerance to environmental stresses, which could be expressed in a hexaploid genetic background, as lots of related QTLs or genes had been mapped to D genome of synthetic wheat (Nkongolo et al., 1991; Innes and Kerber, 1994; Cox and Hatchett, 1994; Thompson and Haak, 1997; Yang et al., 2003; Zhu et al., 2005; Miranda et al., 2006; Ryan et al., 2010; Sohail et al., 2011; Olson et al., 2013a, 2013b; Wan et al., 2015). But the D genome of the first bread wheats were originated from only a small part of *Ae. tauschii* population that cannot possess all the superiorities mentioned above in a few lucky individuals. These must be some other reason to explain more rapid spread of SHW than the tetraploid wheat.

Interestingly, our study shows that hexaploidization enhanced genetic recombination of the ancestral diploid D genome in allohexaploid wheat. And the RF over the whole D genome in SHW increased more than 2 fold than that in diploidy, which does favor to bread wheat in enhancing variation and adaptive evolution by intercrossing with each other among the first hexaploidy individuals of wheat, as more recombination events has the potential to substantially accelerate the development of new varieties by (1) allowing quick assembly of novel beneficial multi-allelic complexes and (2) breaking the linkage among unfavorable genes and fixing desirable haplotypes in fewer generations (Wijnker and de Jong, 2008). This was more efficient than that in a diploid or tetraploid genetic background, while more recombination events could be obtained in hexaploid genetic background.

Our study suggest that the enhanced genetic recombination of the ancestral diploid genome that was caused by hexaploidization could be regarded as another advantage or a new way to increase evolutionary potential of polyploidy, which have been reported as a shaping force in the evolution by taking the advantage of heterosis and gene redundancy yet (Feldman et al., 1997[10]; Ozkan et al., 2001; Shaked et al., 2001; Feldman and Levy, 2005; Comai, 2005; Chen and Ni, 2006; Chen, 2007). More recombination events could generate more gene combination types or haplotypes for natural or artificial selection, resulting in rapid adaptive evolution. Moreover, considering the narrow origination of D genome of the modern common wheat (Wang et al., 2013), more gene alleles from diverse *Ae. tauschii* species in nature can be introduced to the pool of the morden bread wheat by re-synthetizing hexaploidy, which could also reduce the times to generate enough gene combination types of D genome in SHW and benefit to enhancing variation and adaptive evolution in breeding processes.

## Acknowledgements

This study was partially supported by National Natural Science Foundation of China (Grant No. 31401383), Science and Technology Department of Sichuan Province (Grant No. 2017JY0077) and Sichuan Provincial Finance Department (Grant No. 2016NYZ-015&-016&0030&0049). We are very grateful to Prof. Baorong Lu of Fudan University for the critical review of this manuscript.

